# tbg - a new file format for genomic data

**DOI:** 10.1101/2021.03.15.435393

**Authors:** Philipp Schönnenbeck, Tilman Schell, Susanne Gerber, Markus Pfenninger

## Abstract

**Motivation:** The question of determining whether a Single-Nucleotide Polymorphism (SNP) or a variant in general leads to a change in the amino acid sequence of a protein coding gene is often a laborious and time-consuming challenge. Here, we introduce the tbg file format for storing genomic data and tbg-tools, a user-friendly toolbox for the faster analysis of SNPs. The file format stores information for each nucleotide in each gene, allowing to predict which change in the amino acid sequence will be caused by a variant in the nucleotide sequence. Our new tool therefore has the potential to make biological sense of the unprecedented amount of genome-wide genetic variation that researchers currently face.

**Results:** The new tab-separated file for storing the nucleotide data can be easily analyzed and used for a wide variety of biological research. It is also possible to automate some of these analyses using the additional analysis tools from *tbg-tools*

**Availability:** *tbg-tools* is written in Python and allows the installation from the command line. It can be found on https://github.com/Croxa/tbg-tools.

## 1 Introduction

As sequencing methods become faster and cheaper (Kris A. Wetterstrand (2020)), there are ever-increasing amounts of genomic data and new processing possibilities. The analysis of genomic variants, for instance, marks an essential step in many different areas of biomedicine and medical diagnostics and enhances substantial progress in biological research. To determine a species’ genomic diversity, variant calling with different individuals’ sequencing data is utilized. Either reads of single individuals (individual reseq) or reads of whole populations (poolseq) within a species are sequenced and then compared to the reference genome. These variants are often stored in vcf files (Variant Call Format). It is, for example, possible to associate phenotypes to specific SNPs, for studying the effects of climate change on genomes (Pauls *et al* (2012), Pfenninger *et al* (2020)), or to study mutations and changes in the number of genome copies in cancer cells (Koboldt et al., 2012). In this fast growing world of genomics, more and more important and useful new programs and file formats are therefore developed, such as (Li *et al* (2020)) or GIANT (Vandel *et al* (2020)). However, many of these programs serve a particular niche and furthermore, they lack of answering more general questions about the sequenced data, such as e.g. providing information about synonymous and non-synonymous substitutions. However, this information is critical for diagnostics and biological research, since non-synonymous mutations can often explain phenotype variants.

To close this gap in currently existing tools, we developed the user-friendly toolbox *tbg-tools* available on Github as open source. The program offers a built-in toolbox as well as a new data format called *tbg.tsv*. Using the new format, information on specific variants can be accessed quickly and easily using unix command tools like *awk* (Aho *et al* (1987)) or with *tbg-tools* itself. *tbg-tools* offers a simple command-line based handling and can be installed within seconds using *GitHub*. It requires a Python version of 3.6 or higher but no additional modules that might complicate the installation. For analysis *tbg-tools* only need a fasta file with the genome and an associated gff file with the genome’s coding regions. Since these data files are usually created during de-novo genome assembly and -annotation the creation of a *tbg* file can be easily integrated into an already existing analysis pipeline.

## 2 Methods

The new file format *tbg.tsv* is a tab-separated table that can be written as output by *tbg-tools* (Table 1). This human readable *tsv* file is designed for manual editing and analysis by the user. Internally, *tbg-tools* works with the binary *tbg* format, which is designed and optimized with respect to storage space. *tbg-tools* can use this optimized file format for extracting specific positions in *csv*, *vcf* and *sync* files. The output is a modified version of these files with information about the coded amino acid.

**Table 1.** Example for a tbg.tsv file. For each nucleotide of each gene there is one row with twelve columns. These columns carry the following information: **Scaffold** refers to the ID of the sequence on which the current gene is located. **Position** gives the position of the nucleotide on the sequence defined by the scaffold. The position is 1-based, like the gff files. **base in reference** provides the base located at the coordinate in the sequence. **Coding base** is located in the gene. If the gene is on the same strand as the reference sequence, this column is equal to the base in reference column. Otherwise, it is the complementary base. **Triplet position** is the position in the current encoding triplet. Possible positions are therefore 1, 2 or 3, respectively. **Amino acid coded** is the amino acid encoded by the current triplet. **Gene ID** provides the gene-ID of the current gene. If no substitution of the current bases leads to a change of the amino acid, the column **4d** is “True” - otherwise, it is “False”. The last four columns, **to-A**, **to-T**, **to-C** and **to-G** give the amino acids expected when replacing the current base with A, T, C or G.

| scaffold | position | base in reference | coding base | triplet position | amino acid coded | gene-ID | 4ds | to-A | to-T | to-C | to-G |
| --- | --- | --- | --- | --- | --- | --- | --- | --- | --- | --- | --- |
| scaffold001 | 190 | A | A | 1 | M | gene001 | False | M | L | L | V |
| scaffold001 | 191 | T | T | 2 | M | gene001 | False | K | M | T | R |
| scaffold001 | 192 | G | G | 3 | M | gene001 | False | I | I | I | M |
| scaffold001 | 193 | A | A | 1 | K | gene001 | False | K | stop | Q | E |
| scaffold001 | 194 | A | A | 2 | K | gene001 | False | K | I | T | R |
| scaffold001 | 195 | A | A | 3 | K | gene001 | False | K | N | N | K |
| scaffold001 | 196 | C | C | 1 | R | gene001 | False | S | C | R | G |
| scaffold001 | 197 | G | G | 2 | R | gene001 | False | H | L | P | R |

As an example, for the run-time (figure S1) and the maximum RAM usage (figure S2), *tbg-tools* was challenged with the data from the human genome project version GRCh38 using a different amount of threads each time. The given *fasta* file has a size of 3.1 GB, the *gff* file has 1.1 GB and the *vcf* file used has a size of 0.27 GB. *tbg-tools* also has an option for computers with a lower amount of RAM.

While analyzing *vcf* files, a stats file option exists, which provides the user with some additional statistics about their *vcf* file (see Table S1). These statistics include, among other things the number of positions found in coding regions, the number of synonymous and non-synonymous changes on all chromosomes and the number of variants that are not SNPs (Indels). The additional stats files for each individual of a *vcf* file have some further information for each individual.

## 3 Discussion

We expect a broad field of applications to benefit from using the *tbg.tsv* file with minimal manual effort. An example for the analysis of synonymous and non-synonymous mutations using *tbg-tools* is the calculation and interpretation of the *ω* = *K*_*a*_/*K*_*s*_ ratio (also known as the dN/dS ratio) (Motoo Kimura (1977)). Another potential field of application could be the search for specific amino acid changes during medical diagnostics. If any amino acid of the *α*-helix is substituted by a proline, the helix is most often interrupted, and the functionality of the protein is not guaranteed anymore. Therefore, it could become routine part of the diagnostics analyses to search for those specific substitutions in the amino acid sequence that could cause structural changes or a stop codon.

Because the *tbg.tsv* file has a column indicating 4-degenerate sites it can be utilized to identify these positions. 4-degenerate sites can be used e.g. in the calculation of the site frequency spectrum at neutral sites (Chueca *et al* (2021)) and guide the detection of positive selection in coding sequences (Künstner *et al* (2010)). Overall, *tbg-tools* and the new format *tbg.tsv* are versatile for various analyses and can be easily added to existing genome analysis pipelines without consuming much time while still being easy to install and use.

## Acknowledgement

The authors acknowledge funding from the Emergent AI Center funded by the Carl-Zeiss-Stiftung and thank the LOEWE-Centre TBG funded by the Hessen State Ministry of Higher Education, Research and the Arts (HMWK).

