## Supplement for "tbg - a new file format for genomic data"

### tbg - a new file format for genomic data: supplemental document

#### 1. FIGURES AND TABLES

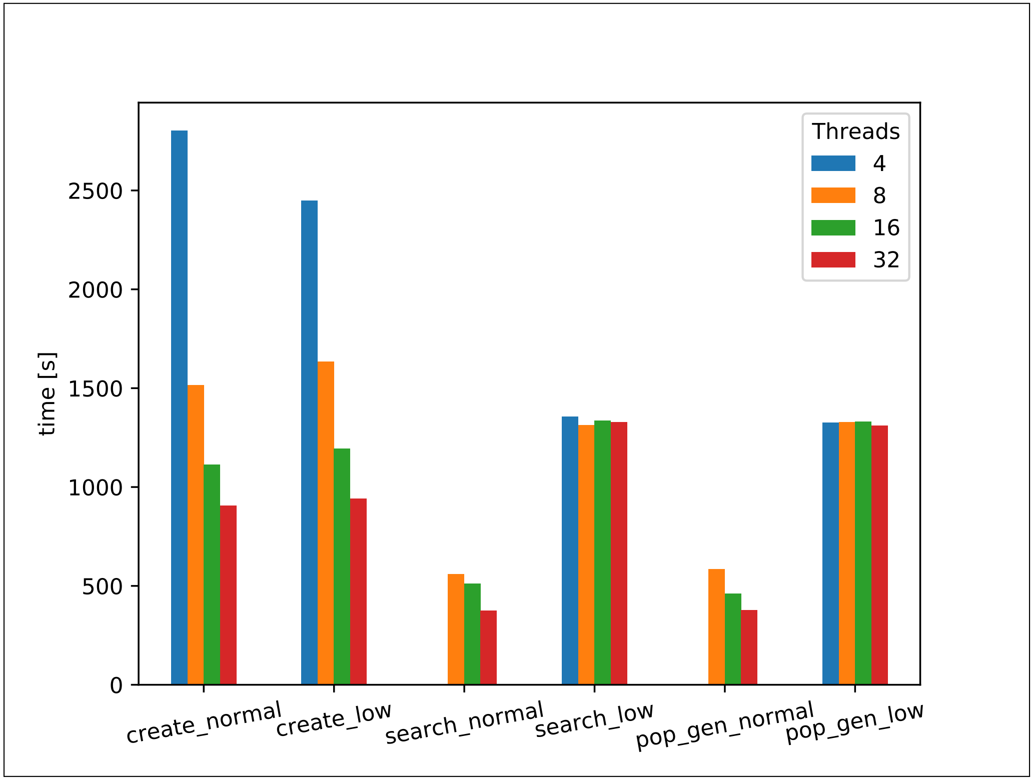


**Fig. S1.** Run-time in seconds [s] of the subprograms *create*, *search* and *pop_gen* of *tbg-tools* with 4 (blue), 8 (orange), 16 (green) and 32 (red) threads. The *low* text represent the usage of the *low RAM* option. Because of built-in limitations in the multiprocessing python package, the number of threads used in search and pop_gen has to be higher than the size of the tbg file in GB. Therefore, there is no data about run-time and RAM usage for these subprograms using four threads because the *tbg* file was larger than 4 GB.

**Table S1. Example of a stats-file of the human genome *vcf* file**

| | |
|---|---|
| SNPs found in coding sequences | 3371756 |
| SNPs not found in coding sequences | 152161 |
| variants which are not SNPs | 373777 |
| synonymous_changes | 1165269 |
| non_synonymous_changes | 2206487 |

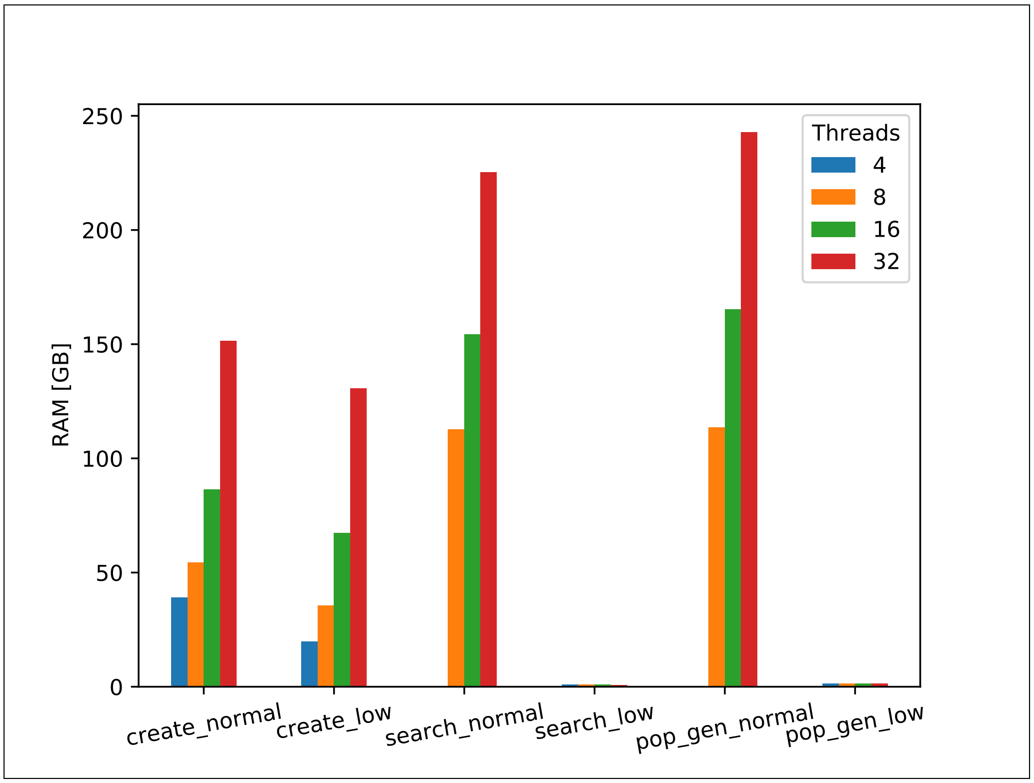


**Fig. S2.** Maximum of RAM usage in gigabytes [GB] of subprograms *create*, *search* and *pop_gen* of *tbg-tools* with 4 (blue), 8 (orange), 16 (green) and 32 (red) threads. The *low* text represent the usage of the *low RAM* option. Because of built-in limitations in the multiprocessing python package, the number of threads used in search and pop_gen has to be higher than the size of the tbg file in GB. Therefore, there is no data about run-time and RAM usage for these subprograms using four threads because the *tbg* file was larger than 4 GB.
